# Determinants of accurate virus assembly: *in vitro* assembly of a complex icosahedral viral capsid

**DOI:** 10.64898/2026.11.26.769398

**Authors:** Franziska Glas, Marius Obst, Katrin Wöhrle, Jochen Reinstein

## Abstract

Understanding how capsids, the protein shells of viruses, assemble is crucial to decipher viral life cycles. However, the assembly process is complex, particularly for larger viruses, and challenging to study in vitro. In this study, combining electron microscopy with biochemical assays, we demonstrate that the capsids of the virophage Mavirus can self-assemble *in vitro* using solely the information encoded within its major capsid protein. Polyanions act as non-specific charge gatherers, increasing local concentrations of capsomers without serving as templates or possessing sequence specificity. Crucially, the identity and length (as long as ≥ 75 bp) of the polyanion - whether DNA or another polyanion - do not affect capsid assembly, underscoring that DNA does not act as a “tape measure”. Furthermore, the unstructured and extended C-terminal region of the major capsid protein also does not govern capsid size. The cryo-EM structures of the Mavirus capsid reveal a triangulation number of T = 27 and exhibit high structural complexity, highlighting the precision and robustness of the self-assembly process.

## Introduction

The capsid, the outer protein shell of a virus particle, encloses and protects the viral genome and contains components that ensure specific docking to host cells. Knowledge of capsid assembly and maturation not only allows us to understand diverse viral life cycles, but also contributes to the development of antiviral compounds as well as nanocontainers with a broad range of practical applications. Capsids are diverse in structure but are always composed of viral capsid proteins (capsomers) as the principal building blocks ^1^; the capsomers assemble into capsids in an oligomerization reaction with highly volatile intermediates ^2^. The assembly of capsids through these intermediates is an extremely delicate and complex process that requires, particularly in the case of larger viruses, a multitude of accessory components ^3^. Thus, the study of the assembly pathway is often challenging and experimentally difficult.

Based on the structure of their capsid proteins, four major lineages of icosahedral viruses have been described: PRD1/adenovirus-like, Picorna-like, HK97-like and BTV-like ^4^. So far, systematic studies of *in vitro* capsid assembly have focused on small viruses whose capsid architecture is characterized through a low triangulation (*T*-) number that refers to the number of similarly structured subunits per asymmetric unit of the icosahedral capsid ^1^ and thus determines complexity of the virus shell. Well-characterized viral assembly systems include in particular cowpea chlorotic mottle virus (CCMV, *T* = 3), hepatitis B virus (HBV, *T* = 4), bacteriophage P22 procapsid (*T* = 7) and simian virus 40 (SV40, *T* = 7). However, the known assembly systems rarely exceed *T* = 7 capsid architectures and include viruses of either the Picorna-like, HK97-like, or BTV-like structure-based lineage but no representatives of the well-established and ubiquitous PRD1/adeno-like lineage. Members of this lineage infect hosts from all three domains of life and possess by far the largest and most complex viral capsids known in nature. Prominent members are adenovirus (*T* = 25) ^5^, PRD1 bacteriophage (*T* = 25) ^6^, viruses of the proposed order *Megavirales* ^7^ with *T*-numbers of up to ∼1,000 and also the virophages Sputnik (*T* = 27) ^8^ and Mavirus ^9^. Capsids of the PRD1/adeno-like lineage share the unique feature that they are composed of two distinct capsid proteins with a jelly-roll fold, referred to as major and minor capsid proteins. In the trimeric major capsid protein (MCP_3_), two consecutive jelly-rolls are linked within a single protomer, resulting in a double jelly-roll fold and pseudo-sixfold symmetry ^10^. A pentameric minor capsid protein (penton) with a single jelly-roll fold forms the capsids’ vertices ^11^. All viral capsids known so far to be capable of assembling *in vitro* consist of only one capsid protein, filling both pentavalent and hexavalent capsid sites ^12–18^. Since viral assembly has mostly been studied using small viruses composed of one capsid protein, it is also important to address the question how larger viruses with two capsid proteins assemble.

Theoretical studies have predicted that capsids with *T* > 4 require templates such as scaffolding proteins or inner cores ^19^. Without a template, large icosahedral shells (*T* > 4) did not assemble in these simulations; but tubular structures were obtained instead. In more recent theoretical studies, however, icosahedral capsids with higher *T*-numbers assembled even in the absence of a template ^20^.

We present here a quantitative experimental study of the assembly of the marine virophage Mavirus, a member of the PRD1/adeno-like lineage. Mavirus-sized virus-like particles (VLPs) have been reported to assemble spontaneously from the trimeric MCP_3_ in *E. coli*, indicating that viral- or host-specific factors are not required ^21^. We used this system to study assembly *in vitro*, determined the overall capsid architecture of *in vitro* assembled Mavirus by cryo-electron microscopy (cryo-EM) reconstructions and analyzed assembly conditions using an online *in vitro* assembly assay based on monitoring static light scattering (SLS). We show that assembly of these larger and more complex Mavirus capsids is polyanion-triggered, but requires neither scaffolding nor tape measuring, in line with recent theoretical predictions by Mendoza and Reguera ^20^. The polyanion serves merely to enhance the local capsid protein concentration. Our findings indicate that neither the length of the C-terminal domain nor the presence of double-stranded DNA governs capsid size. Instead, the precise determination of capsid size and the establishment of icosahedral symmetry are exclusively encoded within the core interface of the major capsid protein.

## Results

### “Whiffle balls” assemble during MCP_3_ purification

Mavirus-sized virus-like particles self-assemble to capsids from MCP_3_ capsomers when expressed in *E. coli* ^21^. In order to obtain assembly competent capsomers for in vitro assembly studies, we added the N-terminal purification tag, maltose binding protein (MBP), which prevented MCP_3_ from assembling to capsids. Removal of the tag by proteolytic cleavage during purification induced part of the untagged capsomer to assemble *in vitro*. Negative-stain electron microscopy showed VLPs, indicating that assembly from MCP_3_ is possible *in vitro* (**Fig. 1B**). Since these particles lack the pentameric penton protein at the capsids’ vertices, they were termed “whiffle balls”, in analogy to particles described for a mutant of the virus HK97 ^22^. Mavirus whiffle balls mostly appeared as spherical particles in negative stained electron microscopy samples and the existence of an equilibrium between free MCP_3_ and capsids could be observed (**Fig. S1B**). Moreover, subsequent incorporation of penton at the vertices rendered the whiffle balls icosahedral (**Fig. 1**).

**Fig. 1.**
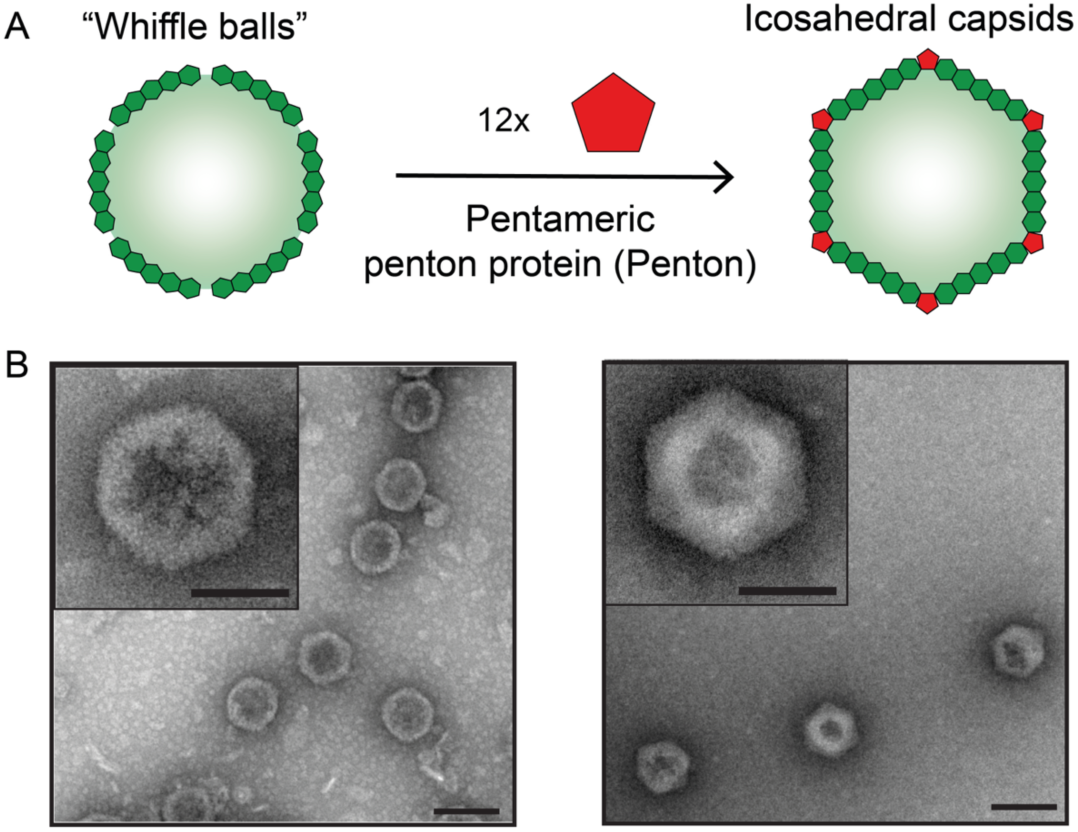
*In vitro* assembly of *T*=27 Mavirus capsids. (**A**) Sketch of Penton incorporation. Spherical whiffle balls are solely assembled of MCP_3_ (green hexagons) with gaps at the 12 capsids’ vertices. Adding the pentameric penton protein (red pentagons) to *in vitro* assembled whiffle balls renders them icosahedral. (**B**) Negative-stain TEM of purified whiffle balls assembled during MCP_3_ purification (left) and after penton incorporation (right). The insets show a close-up of the particles. Scale bars represent 50 nm and 100 nm, respectively.

### Polyanions trigger Mavirus *in vitro* capsid assembly

The C-terminal domain (CTD) of Mavirus MCP is highly positively charged (theoretical isoelectric point: 10.15), which suggests that electrostatic interactions with nucleic acids are a significant part of the thermodynamic driving force for capsid assembly *in vivo*. Capsids of both RNA and DNA viruses have been assembled *in vitro* around heterologous nucleic acids and negatively charged non-nucleic acid compounds, indicating that sequence specificity is in many cases not required ^23–25^. Empty viral capsids have been assembled *in vitro*, without the need for polyanions, by increasing ionic strength ^17,26–28^, lowering pH ^29,30^ or even removing reducing agents to form disulfide crosslinks between capsomers ^31^. To probe the assembly process under more physiological conditions we tested the effects of polyanions including polystyrene sulfonates (PSS), poly-γ-glutamic acids (γ-PGA) and nucleic acids of unspecific sequence on the assembly of Mavirus capsids *in vitro*. Complete whiffle balls assembled irrespective of the type of polyanion and without the need for any additional scaffold protein or tape measuring under physiological conditions (**Fig. S1**) as long as MCP_3_ were assembly competent – only unassembled MCP_3_, which eluted at 28.8 – 32.5 mS/cm during cation exchange chromatography (IEX), were assembly competent (see methods). Hence, we propose that during protein expression and purification in *E. coli* small nucleic acid fragments or other polyanionic compounds such as glycosaminoglycans (*e.g.* hyaluronic acid, heparin, dextran sulfate) were attached to the CTD, which in turn prevented polyanion interaction and capsid assembly. These MCP_3_ were assembly-incompetent as well as those that lost the C-terminus. It is also suggested that the negatively charged fragments or compounds attached to the CTD might be partially competed off using IEX and rendered assembly-incompetent MCP_3_ to their assembly-competent form. This observation could be relevant for all viral systems where similar charges are present and capsid assembly is triggered by electrostatic interactions of the general capsid building blocks and polyanions.

### Capsid size depends solely of capsomers

We further investigated whether the size of the capsid (whiffle ball) could be controlled by either the length of a polyanion, in this case dsDNA or the C-terminal domain (CTD). Remarkably, capsid size remained constant across all DNA lengths tested, indicating that DNA does not dictate capsid dimensions (**Fig. S2**). However, DNA length significantly influenced both assembly efficiency and stability. With DNA lengths of 50 bp whiffle balls assembled inefficiently (**Fig. S2A**), whereas DNA lengths of 75 bp or greater exhibited efficient assembly and improved stability, as reflected in the assembly curves and electron micrographs. As DNA length increased, the yield of intact whiffle balls decreased. At 3,000 bp and especially at 20,000 bp, the equilibrium shifted away from capsids, favoring free major capsid protein (MCP_3_) or the formation of non-capsid structures such as tubular assemblies (**Fig. S2B**). This demonstrates that while DNA length affects assembly efficiency, it does not influence capsid size - highlighting a fundamental uncoupling between structural scaffolding and size determination in Mavirus capsids.

To assess the role of the CTD in capsid assembly and size determination, we generated eight Mavirus capsid protein mutants: four with truncated CTDs (–15, –30, -45, –60 residues) and four with extended CTDs (+15, +30, +45, +60 residues). Surprisingly, capsid size remained unchanged across all variants and the wild type as control —except the –60 mutant, which failed to assemble entirely (**Fig.2, S3**). For the remaining mutants, capsid diameters were consistent and showed minimal variation (measured across up to 170 particles per variant, with small measurement errors). Data are summarized in Fig. 2, confirming that the CTD does not regulate capsid size, unless it is severely truncated.

**Fig. 2.**
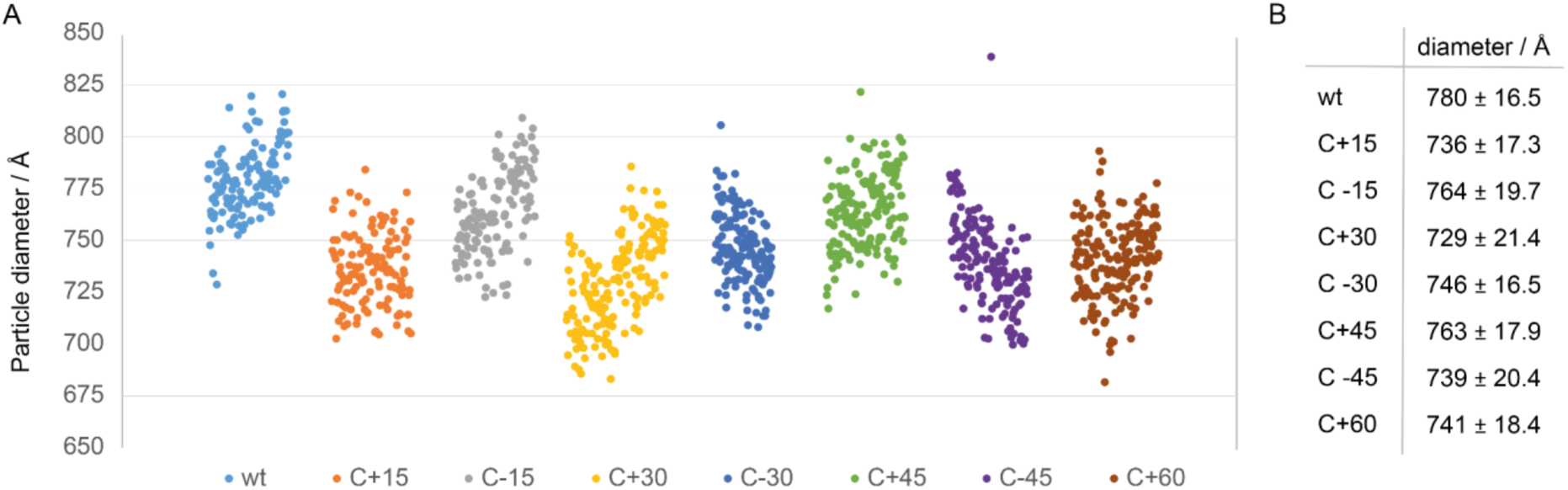
Measurement of whiffle ball diameter after assembly with varying DNA length. (**A**) Diagram of the statistical analysis of the whiffle ball particle diameter of each variant. The parameters were obtained from 130-170 individual particles. (**B**) The mean value ± the standard deviation of the particle measurement study.

### Electron cryo-microscopy

After optimizing and identifying stable conditions for the VLP assembly, electron cryo-microscopy was performed of purified whiffle balls and whiffle balls with incorporated penton (VLP) to determine the overall capsid composition and the symmetry and complexity of Mavirus capsids. We used single-particle analysis to process the data yielding resolutions of 4.2 Å for the whiffle ball and 5.2 Å for the complex with bound penton (**Fig. S.4**). The cryo-EM density maps and the crystal structures of MCP_3_ (PDB-ID: 6G45) and the penton protein (PDB-ID: 6G42) allowed us to build atomic models of the complexes (**Fig. 3A,B**). Each structure comprises 260 major capsid proteins (MCP_3_) and in addition 12 pentons in the VLP structure. Both structures exhibit T = 27 quasi-symmetry (**Fig. 3A**, h = 3, k = 3), similar to the Sputnik virophage ^8^. As reported for Sputnik, the icosahedral asymmetric unit (IAU) of Mavirus was also shown to consist of 4 central pseudohexameric MCP_3_, one third of a pseudohexameric MCP_3_ positioned at the icosahedral 3-fold axis, and one fifth of a pentameric penton at the 5-fold axis (**Fig. 3C**). Interestingly, one pseudohexameric MCP_3_ is rotated in the Mavirus structures compared to Sputnik. We investigated how the inter-subunit interface in Mavirus VLPs is affected by rotation of the MCP_3_ protomer at the icosahedral 3-fold axis. Upon rotation, one of the four interfaces with subunit I is completely disrupted (**Fig. 3D**), suggesting that this conformational change enhances VLP stability by reorganizing MCP_3_-MCP_3_ contacts. Increased interface connectivity correlates with greater structural stability. However, the intrinsic interactions between individual MCP_3_ subunits are weak: no significant hydrophobic or polar interaction networks are present. Only 1 - 2 hydrogen bonds and single salt bridges stabilize adjacent protomers. This limited interaction strength is consistent with the observed dynamic behavior of the VLP, which exhibits flexibility reminiscent of a “whiffle ball,” reflecting a metastable, adaptable architecture. Variability was also observed in the cryo-EM data of the whiffle ball. The 2D class averages (**Fig. S4G**) showed both icosahedral and spherical particles, suggesting that the sample might exists in an equilibrium between the icosahedral (whiffle ball) conformation, which is competent for penton binding, and a more compact spherical state. This conformational heterogeneity is consistent with relatively weak intrinsic interactions.

**Fig. 3.**
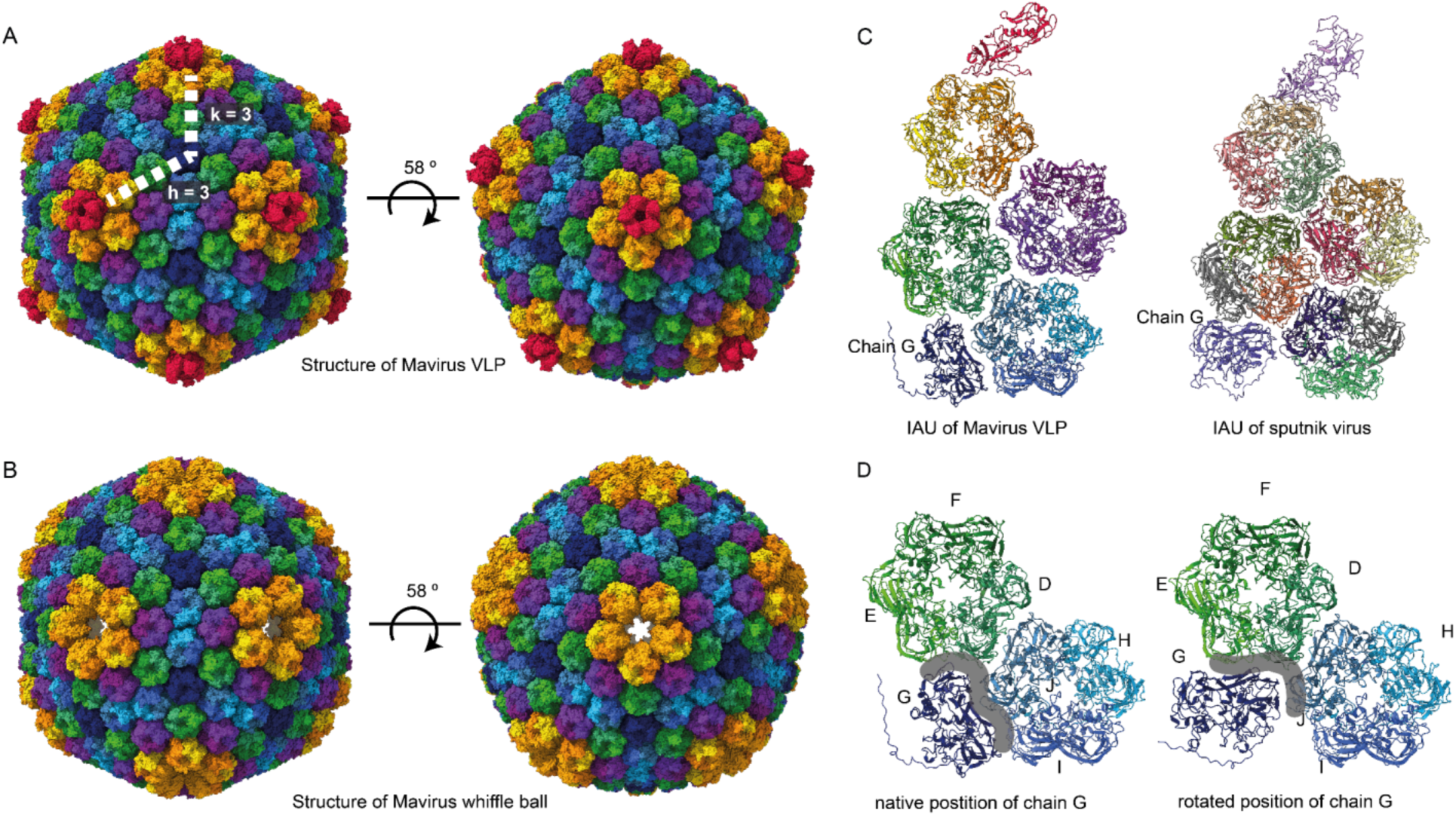
Structural overview of Mavirus VLP and whiffle ball. (**A**) Surface view of the capsid structure resolved with cryo-EM at 5.2 Å. The capsid is composed of 260 MCP_3_ subunits and 12 penton proteins (red). The identification of the triangulation number of *T*=27 (*h*=3, *k*=3) is indicated. (**B**) Surface view of the whiffle ball structure resolved with cryo-EM at 4.2 Å, lacking the penton protein. (**C**) Icosahedral Asymmetric Unit of Mavirus VLP and asymmetric unit of sputnik (PDB-ID: 3j26). The rotated MCP chain G is annotated. (**D**) Close-up view of the interface of Mavirus VLP and the rotation of the chain G. The interface between chain G and the other subunits is indicated with the grey area.

Overall structural comparison reveals only minor differences between the whiffle ball and the VLP. Notably, in the icosahedral whiffle ball the penton-binding pocket is already pre-formed, indicating that no major conformational rearrangement is required for penton binding - only a slight increase in local radius (**Fig. 4**). Upon penton binding, the pocket expands by approximately 4 Å (**Fig. 4C**). Given the symmetric arrangement of 12 pentons, this local expansion is amplified across the capsid, resulting in an overall increase in diameter of about 50 Å. This predicted structural enlargement is consistent with experimental observations showing that the complete VLP with bound pentons is slightly larger than the whiffle ball alone (diameter increases from 713 to 776 Å, **Fig. 4D**), supporting a model of cooperative, symmetric structural adaptation during penton incorporation.

**Fig. 4.**
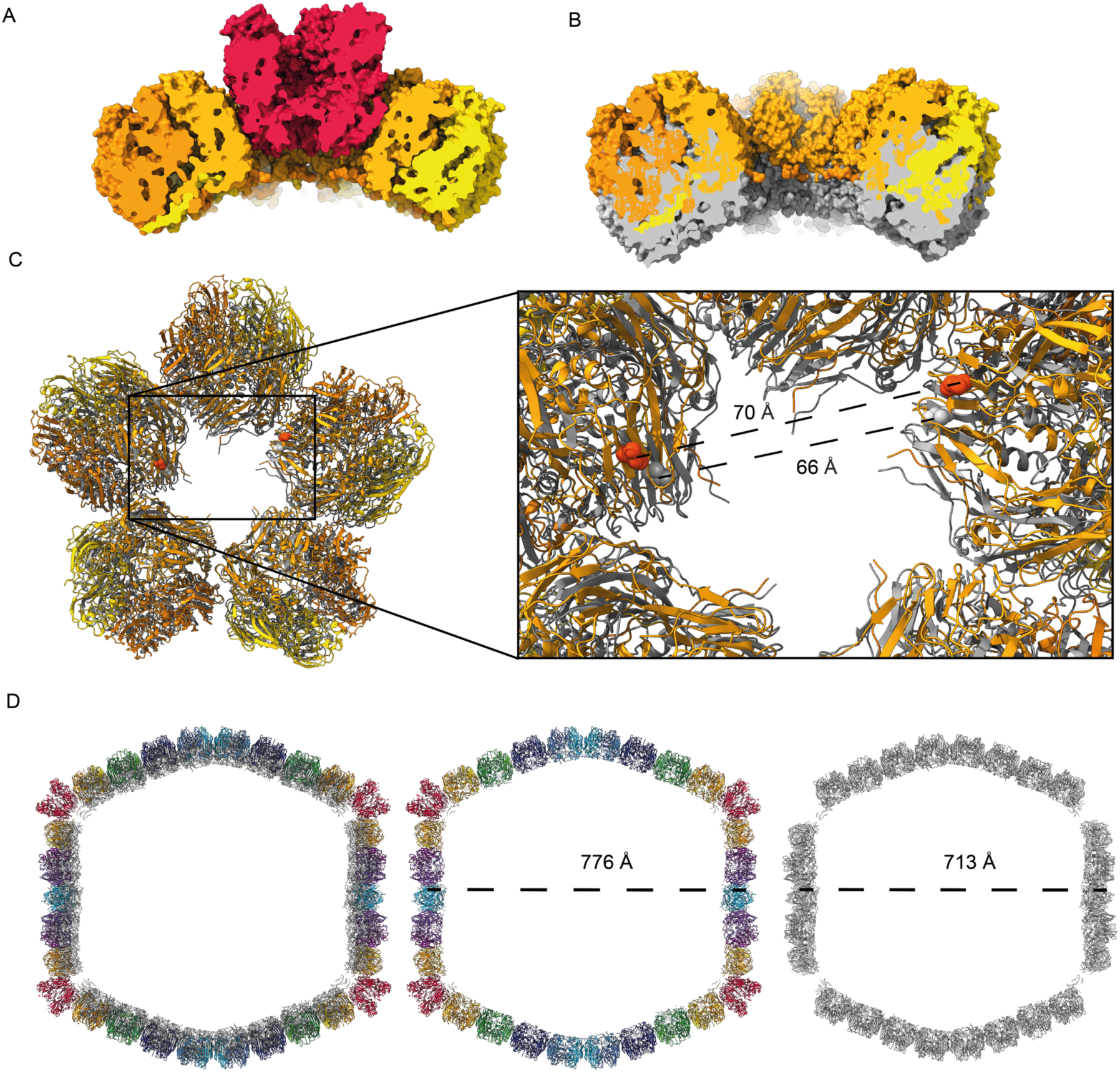
Penton binding region of Mavirus VLP. (**A**) Cross-sectional view of the VLP surface structure, highlighting the penton-binding interface. (**B**) Overlay of whiffle ball (grey) and VLP, focusing only on the MCP_3_ subunits involved in penton binding. (**C**) Structural overlay of the five MCP3 subunits forming the penton-binding region in VLP (orange-yellow) and whiffle ball (grey). Two Ile 227 residues are highlighted. The right panel shows a zoomed view of the Ile 227 side chains, revealing a reduced inter-residue distance in whiffle ball compared to VLP. (**D**) Detailed view of a single MCP_3_ ring, illustrating the diameter differences between the two variants. Left: overlay of VLP and whiffle ball; middle: VLP alone; right: whiffle ball alone

## Discussion

We demonstrate here that *in vitro* assembly of a *T* = 27 viral capsid of the PRD1/Adeno family can occur without additional scaffolding proteins and tape measuring. We showed that despite its large size (d=70 nm), complex architecture, capsids of uniform size assemble reliably under defined conditions. All information is encoded solely in its trimeric major capsid protein and assembly can be triggered by polyanions through enhanced proximity of the capsomer units. Polyanions are important for charge neutralization and for achieving a critical local concentration of capsomers needed specifically in the case of weak inter-capsomer interactions. We established optimal conditions for Mavirus capsid assembly in order to use Mavirus as a new and appropriate viral assembly system for dsDNA viruses of the PRD1/adeno-like lineage. Based on simulations of virus assembly around nucleic acids, two mechanisms were proposed that depend on the strength of subunit–subunit and subunit–polyelectrolyte interactions ^32^. We propose an *en masse* mechanism with strong subunit–polyelectrolyte interactions but rather weak subunit–subunit interactions. This is also consistent with the fact that we could not assemble capsids with the capsomer alone.

Using this system, we showed that Mavirus assembly *in vitro* using highly variable nucleic acid lengths produces nevertheless capsids of the same size and morphology, as long as the provided DNA length is above about 50 bp. It still needs to be elucidated whether capsids are also able to assemble around the full dsDNA genome of Mavirus (∼19 kbp), which would address the biological implication of Mavirus assembly in more detail. In this case one would have to ask why all virophage genomes contain the sequence of a highly conserved Ftsk-type ATPase predicted to be responsible for active DNA transport ^33^.

Nevertheless, judging from our results, *in vivo* assembly should occur around any nucleic acid present making it unlikely for polyanions to function as a true assembly template. They mainly serve to gather charges and thus bring capsomers into close proximity, as also seen for viruses of other lineages ^25,34,35^ but at least in the case of Mavirus do not determine size. If a structured assembly platform is required to determine size and shape, as has been predicted to be necessary for all *T* > 4 viral capsids ^19^, only scaffolding proteins or inner cores such as lipid membranes or smaller capsid structures could provide this. Since no additional components were present in the assembly reaction (except DNA or other polyanions) the only remaining candidate as a potential scaffold is the unstructured highly positive C-terminus of a about 100 aa residues. It should be noted that this part of the capsomer is not visible in the X-ray nor the EM densities. Our data presented here with constructs that largely varies in length of the C-terminus indicate that an intrinsic scaffold protein can also be ruled out as assembly size determinant.

The present results expand our knowledge of how larger viral capsids assemble and suggest that capsids which include two distinct capsid proteins, as is the case only for viruses of the PRD1/adeno-like lineage, might in general also carry information on size and shape in the major capsid protein. However, *in vivo* ordered assembly platforms composed of appropriate scaffold proteins or inner cores may still be required to increase local capsomer concentrations when nucleic acid interactions are absent.

As one of the smallest and least complex representatives of the PRD1/adeno-like lineage, Mavirus capsids represent a promising platform for the development of viral nanocontainers. As a first step, Mavirus capsids assembled around designed DNA fragments could serve as a natural DNA delivery system, initially targeting their native host, *Cafeteria roenbergensis*.

## Materials and Methods

### Purification of Mavirus capsid proteins and Virus Like Particles (VLPs)

Full-length and C-terminal truncated MCP_3_ was expressed and purified using an N-terminal purification tag (comprising maltose binding protein (MBP) and His_6_) as previously described ^21^, except for using 50 mM HEPES-NaOH, 10 mM NaCl, 1 mM TCEP, pH 7.0 as dialysis buffer for cation exchange chromatography (Resource-S) and 50 mM HEPES-NAOH, 1 mM TCEP, 150 mM NaCl, pH 7.0 for size exclusion chromatography (SEC). Assembly-competent MCP_3_ eluted at 28.8 – 32.5 mS/cm during cation exchange chromatography and were further purified with SEC chromatography to remove minor fractions of VLPs that were forming through the desalting effect of the SEC run.

VLPs could be produced upon addition of polyanions to assembly competent MCP_3_ obtained as described above, but were also harvested during purification of full-length MCP_3_ in *E. coli*. In brief, VLPs assembled after TEV protease cleavage of the N-terminal MBP-purification tag and were separated from unassembled MCP_3_ using Ni^2+^-affinity chromatography by following the A260/A280 ratio. VLPs (A260/A280 > 0.8) were subjected to SEC using a Superdex 200 16/60 column (GE Healthcare) equilibrated in 50 mM HEPES-NaOH, 150 mM NaCl, pH 7.0 and VLP containing fractions were pooled, concentrated, and stored at -80 °C. Protein concentration was determined according Ehresmann *et al.* ^36^ since this approach approximated protein concentration of VLPs with sufficient accuracy. A direct determination through A280 and the given extinction coefficient is highly error prone due to the strong light scatter of the 70 nm VLP particles ^37^.

The minor capsid protein (penton) was expressed and purified as described ^21^.

### Cloning of C-terminal variants

The insert plasmids pEX-A128-Mfel_C1±15/-30/-45/-60 and pEX-A258-Mfel_C+30/+45/+60 with the respective genes for various C-terminal MCPs were obtained from eurofins. The vector pET-1a-MBP-MV18_casette for insertion was obtained from Genescript. The sequences of final constructs are listet in the supplements. The insert plasmid and the vector (pET-1a-MBP-MV18_casette) were digested, using the Type IIS restriction enzymes 5’ MfeI, 3‘ HindIII. After ligation, the successful incorporation of the insert was verified with sequencing. Expression and purification as described previously.

### Nucleic acid fragments and other polyanions

For inducing *in vitro* capsid assembly of Mavirus MCP_3_, the following polyanions were screened: poly(4-styrenesulfonic acid) 70 kDa, 200 kDa, and 1 MDa (PSS, Sigma-Aldrich) and low-molecular and high-molecular poly-γ-glutamic acid (γ-PGA.Na, 200-400 kD and 2 MD, Molecular Dimensions). Unless indicated otherwise, γ-PGA refers to the low-molecular form. The 100 bp dsDNA fragment was obtained by annealing complementary Mavirus Terminal Inverted Repeat (TIR) oligonucleotides at 96 °C for 5 min followed by slowly cooling to RT. Annealing was verified by agarose gel electrophoresis. All digestions were performed according to the manufacturer’s instructions and fragments were purified by standard DNA clean up procedures (Promega). For studying the capsid assembly with varying DNA length, dsDNA fragments were obtained from Thermo Fisher Scientific (NoLimits DNA of 75, 250, 500, 1000, 3000 and 20000 bp).

### Online *in vitro* capsid assembly assay

The assay was carried out in a 96-well quartz microplate (Hellma) using the TECAN Spark multimode microplate reader with injector module. The assay is designed in order to screen assembly conditions in a higher throughput. Full-length MCP_3_ was added manually at the respective concentration in the respective assembly buffer and the baseline was measured. When possible, polyanion solution was injected automatically in order to initiate the assembly reaction. For all experiments, a humidity cassette was used to prevent sample evaporation and shaking steps were applied (orbital, 150 rpm, 4 mm – after polyanion addition and before each measurement). Unless indicated otherwise, the assay was performed at 20 °C. In absorbance mode, the apparent changes of static light scattering (SLS) were monitored at λ_1_ = 250 nm when working with non-absorbent polyanions such as γ-PGA or λ_2_ = 310 nm when working with nucleic acids (10 flashes, settle time 50 ms) in order to maximize the scatter signal for low protein concentrations.

### Capsid assembly with varying DNA length and C-terminal variants

The *in vitro* capsid assembly assay using the TECAN Spark multimode microplate reader was performed for analyzing the influence of DNA length on the Capsid assembly. Assembly conditions were 1 μM of MCP_3_ and 15 ng/μl of the respective NoLimits DNA (Thermo Fisher Scientific) in 60 mM UB-NaOH, 150 mM NaCl, pH 7.0. Samples were further analyzed with electron microscopy.

For the C-terminal variants, 1 μM of the respective MCP_3_ mutant and 30 ng/μl of the 100 bp dsDNA TR were assembled in in 60 mM UB-NaOH, 150 mM NaCl, pH 7.0 using the TECAN Spark multimode microplate reader. The statistical size analysis of the C-terminal variants was carried out 4 h after the *in vitro* capsid assembly. The diameter of the particles was measured using the EMAN2^38^ software package. Around 40 micrographs per mutant were taken to obtain 130 – 170 particles for the statistical analysis and to calculate the mean and standard deviation. The measured data is in the supplement.

### *In vitro* assembly for electron microscopy studies

Mavirus major capsid protein (MCP_3_) was mixed with 100 bp dsDNA at a 1:5 molar ratio (5 μM DNA:25 μM MCP_3_) in assembly buffer (50 mM HEPES–NaOH, 150 mM NaCl, pH 7.0) at RT for at least 1 h or at 4 °C overnight. Following incubation, samples were centrifuged at 8,000 × *g*, 4 °C, for 10 min to remove aggregates or large debris. The supernatant was loaded onto a Superdex 200 10/300 increase size-exclusion chromatography column equilibrated in the same buffer. Fractions corresponding to the capsid peak were pooled and concentrated using Amicon® centrifugal filters (30 kDa MWCO) at 5,000 × *g*, 4 °C, for 2 min. The fresh sample was directly transferred on a negative stain or cryo-EM grid.

### Penton incorporation

*In vitro* assembled whiffle balls with 100 bp DNA stored in storage buffer (pH 7.0) were mixed with an equal volume of incorporation buffer (60 mM UB-NaOH, 150 mM NaCl, pH 5.0). The required penton concentration was determined based on the MCP_3_ concentration: [MCP_3_] × 12 ÷ 260 = [penton]. Penton was added in 5-times excess and incubated over night at 4 °C. The sample was then loaded onto a Superdex 200 10/300 increase size-exclusion chromatography column equilibrated in 50 mM HEPES–NaOH, 150 mM NaCl, 1 mM TCEP, pH 7.0. Fractions corresponding to the capsid peak were pooled, concentrated using an Amicon® Ultra-0.5 mL 50 kDa MWCO centrifugal filter at 5,000 × *g*, 4 °C, for 2 min and were directly used for EM studies.

### Sample characterization by negative-stain TEM

Glow discharged formvar/carbon-coated 100 mesh Cu grids were incubated with 4 μl sample diluted to 0.17 μM MCP_3_. Excess of liquid was removed and grids were incubated with 10 μl of 1 % paraformaldehyde (PFA). Grids were washed three times with ddH_2_O and stained with 0.5 % uranyl acetate for 90 s. TEM grids for the analysis of the C-terminal variants and the different assembly approaches with varying DNA size were stained on glow discharged formvar/carbon-coated 100 mesh Cu grids with 4 μl sample diluted to 3 μM MCP_3_. Excess of liquid was removed and grids were washed two times with ddH_2_O and stained with 10 μl of 0.5 % uranyl acetate for 90 s. Imaging was performed on a Tecnai T20 electron microscope using 200 kV for acceleration.

### Cryo-EM data collection, processing and reconstruction

Freshly glow-discharged Quantifoil 2/1 grids with additional carbon were used for Mavirus capsid (VLP) and whiffle ball vitrification. The samples were flash-frozen in liquid ethane using a Vitrobot (Thermo Fisher Scientific) at 4 °C and 100 % humidity. 3 μl of a capsid solution (A280 = 4.8) in 60 mM UB-NaOH, 150 mM NaCl, 1 mM EDTA, pH 7.0 was applied on the grid and blotted for 4 s before plunging.

The datasets were collected on a Titan Krios transmission electron microscope (Thermo Fisher Scientific) equipped with a spherical-aberration corrector and an X-FEG (field emission gun). The datasets were collected at the Max Planck Institute of Molecular Physiology, Dortmund. The images were recorded on a Falcon III direct electron detector for VLP and on a K3 for whiffle ball using the automated data collection software EPU (FEI). Four images were acquired per grid hole in all cases. Detailed parameters of data acquisition (total dose and number of frames) are listed in table 1. Frame alignment was performed using MotionCor2 ^39^. In total, unweighted full-dose images, dose-weighted images with full-dose and low-dose were generated. Data processing was performed with the software package Relion 3.1 ^40,41^. Contrast Transfer Function (CTF) estimation was performed with CTFFIND4 ^42^ implemented in Relion3.1 on the unweighted full-dose images. CrYOLO was used for automated particle picking ^43^. Fifteen images were manually picked and used as training data for crYOLO. Particles were extracted from the dose-weighted full-dose images. Initial and final particle numbers are listed in table 1.

Particles were further subjected to 2D classification and 3D classification in Relion to exclude damaged particles or contaminations. We used a low-resolution EM map of the Mavirus VLP filtered to 40 Å as an initial model that we obtained inhouse. For both structures, 3D refinement and 3D classification were performed with imposed icosahedral symmetry. To improve the resolution, particles were CTF-refined and after the final refinement, post-processing was performed to determine the resolution at 0.143 Fourier shell correlation (FSC) criterion (Fig. S4 C,H). Detailed parameters for each dataset are summarized in table 1.

### Model building

The major capsid protein MCP_3_ (PDB-ID: 6G45) and penton (PDB-ID: 6G42) crystal structure were used as starting point for building the atomic models and were initially fitted in the EM density map using rigid body fitting in ChimeraX ^44^. Some regions showed already a good fit and some regions still had to be built or remodeled de novo in Coot ^45^. For de novo model building, residues were only built when we could trace the backbone within the density; otherwise, we deleted these residues. We used the real-space refinement tool of PHENIX ^46^ to refine the models. The geometries of the final refined models were evaluated with MolProbity, and the data statistics are summarized in table 1.

## Supplementary Materials

**Fig. S1. Negative-stain TEM of Mavirus *in vitro* assembly.**

**Fig. S2. Dependence of capsid assembly kinetics and size on DNA length.**

**Fig. S3. Dependence of capsid size on C-terminal length.**

**Fig. S4. EM analysis of Mavirus VLP (A-E) and whiffle ball (F-J).**

**Table 1. Cryo-EM data collection and model refinement statistics Sequences of final constructs of C-terminal variants**.

## Acknowledgments

We would like to thank S. Zimmermann and S. Eisel for excellent technical support and U. Mersdorf for initial help with the cryo-EM samples. We further thank I. Schlichting for intense discussions and A. Christiansen and L. Schmidt for initial work on the major capsid protein. We would like to extend our special thanks to S. Raunser from the MPI of Molecular Physiology in Dortmund for enabling us to collect the EM data of the Mavirus VLPs. We are grateful to O. Hofnagel and D. Prumbaum for performing the data acquisition.

## Funding

This work was funded by the Max Planck Society.

## Author contributions

M.O., F.G., K.W. and J.R. designed and performed research, analyzed and interpreted data. M.O., F.G., and J.R. wrote the paper with input from all authors.

## Competing interests

The authors declare no competing interests.

## Data and materials availability

All data needed to evaluate the conclusions in the paper are present in the paper and/or the Supplementary Materials. The cryo-EM densities of Mavirus capsids VLP and whiffle ball have been deposited in the Electron Microscopy Data Bank under accession nos. EMD-100002 for the whiffle ball and EMD-59128 for the VLP. The corresponding coordinates have been deposited in the Protein Data Bank under accession nos. and PDB-ID 34LA for the whiffle ball and PDB-ID 32RT for the VLP. Correspondence and requests for materials should be addressed to F.G. The expression plasmids for MV17 and MV18 can be obtain through Addgene with the accession codes 249261 (pET-1a-MBP-MV18co) and 249262 (pET-1a-MBP-MV17co), respectively.

## Notes

### Competing Interest Statement

The authors have declared no competing interest.

