## Supplemental data for "Determinants of accurate virus assembly: *in vitro* assembly of a complex icosahedral viral capsid"


**This PDF file includes:**


**Fig. S1. Negative-stain TEM of Mavirus *in vitro* assembly.**

**Fig. S2. Dependence of capsid assembly kinetics and size on DNA length.**

**Fig. S3. Dependence of capsid size on C-terminal length.**

**Fig. S4. EM analysis of Mavirus VLP (A-E) and whiffle ball (F-J).**

**Table 1. Cryo-EM data collection and model refinement statistics**

**Sequences of final constructs of C-terminal variants**

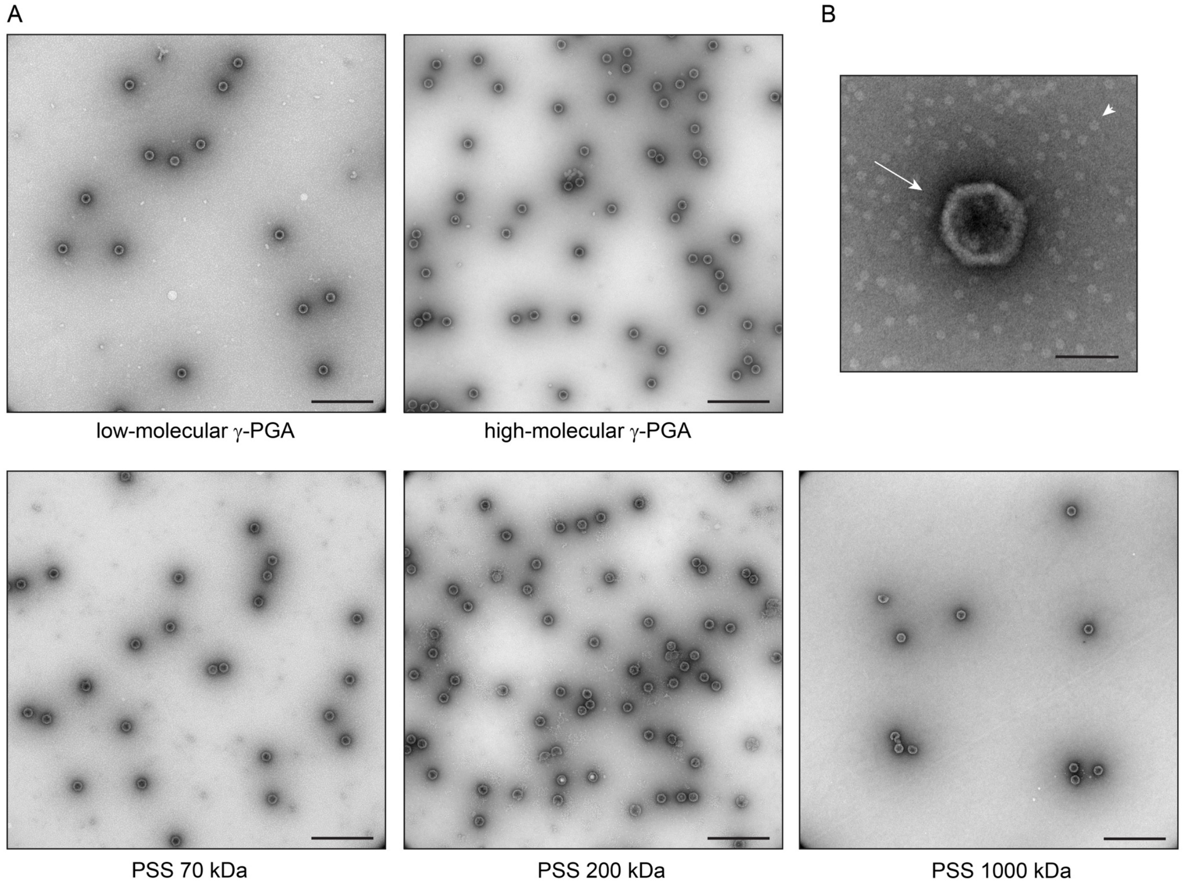


**Fig. S1. Negative-stain TEM of Mavirus *in vitro* assembly.** (**A**) Capsid assembly triggered by various polyanions. Assembly was performed with 1.67 mM $MCP_3$ and polyanion concentrations ranging from 15 to 66 ng/ml at near physiological conditions (150 mM NaCl, pH 7.0). Prior to EM, samples were centrifuged to remove any high molecular assemblies. Scale bars represent 500 nm. g-PGA: poly-g-glutamic acid, PSS: poly(4-styrenesulfonic acid). (**B**) Equilibrium between $MCP_3$ and capsids at 60 mM UB-NaOH, 150 mM NaCl, pH 7.0. The exemplary micrograph of an assembly reaction with $MCP_3$ and low-molecular g-PGA visualizes the equilibrium between unassembled $MCP_3$ (arrowhead) and capsids (arrow). Scale bar refers to 100 nm.

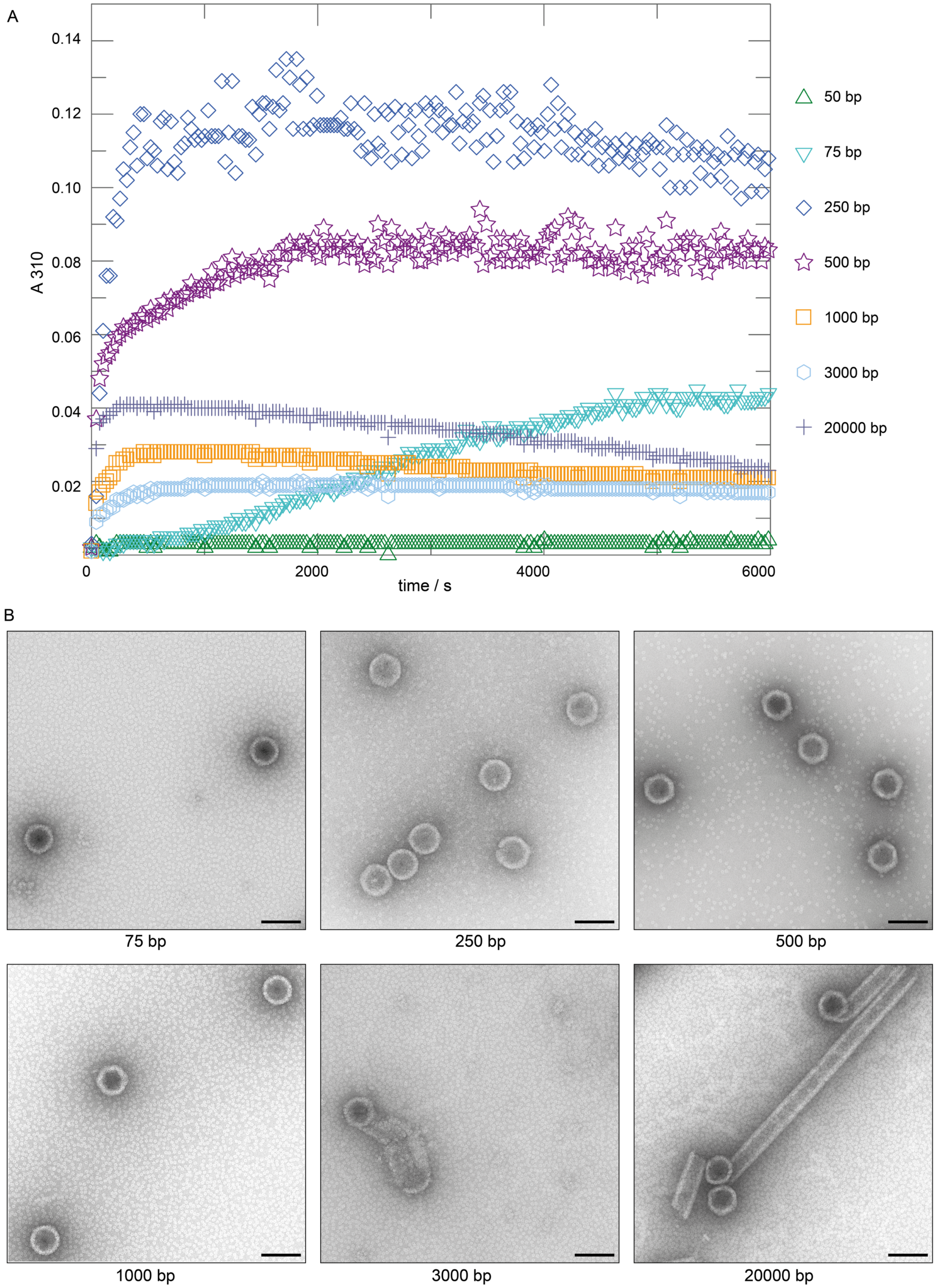


**Fig. S2. Dependence of capsid assembly kinetics and size on DNA length.** (A) MCP capsomers were triggered to assemble by mixing 1 μM $MCP_3$ and 15 ng/ml DNA in 60 mM UB, 150 mM NaCl, pH 7.0. The DNA sizes used were of 50, 75, 250, 500, 1000, 3000 and

20000 bp DNA (Nolimits DNA,Thermo Fisher Scientific). The time course of assembly was followed in a TECAN MTP-reader instrument by following the static light scatter signal at 310 nm. It appears that 50 bp are not sufficient to produce capsids under the given conditions. (**B**) Negative-stain TEM of icosahedral-like whiffle balls assembled as described above. The images represent typical capsids of each sample with unassembled capsomers in the background and some tube structures. Scale bars represent 100 nm.

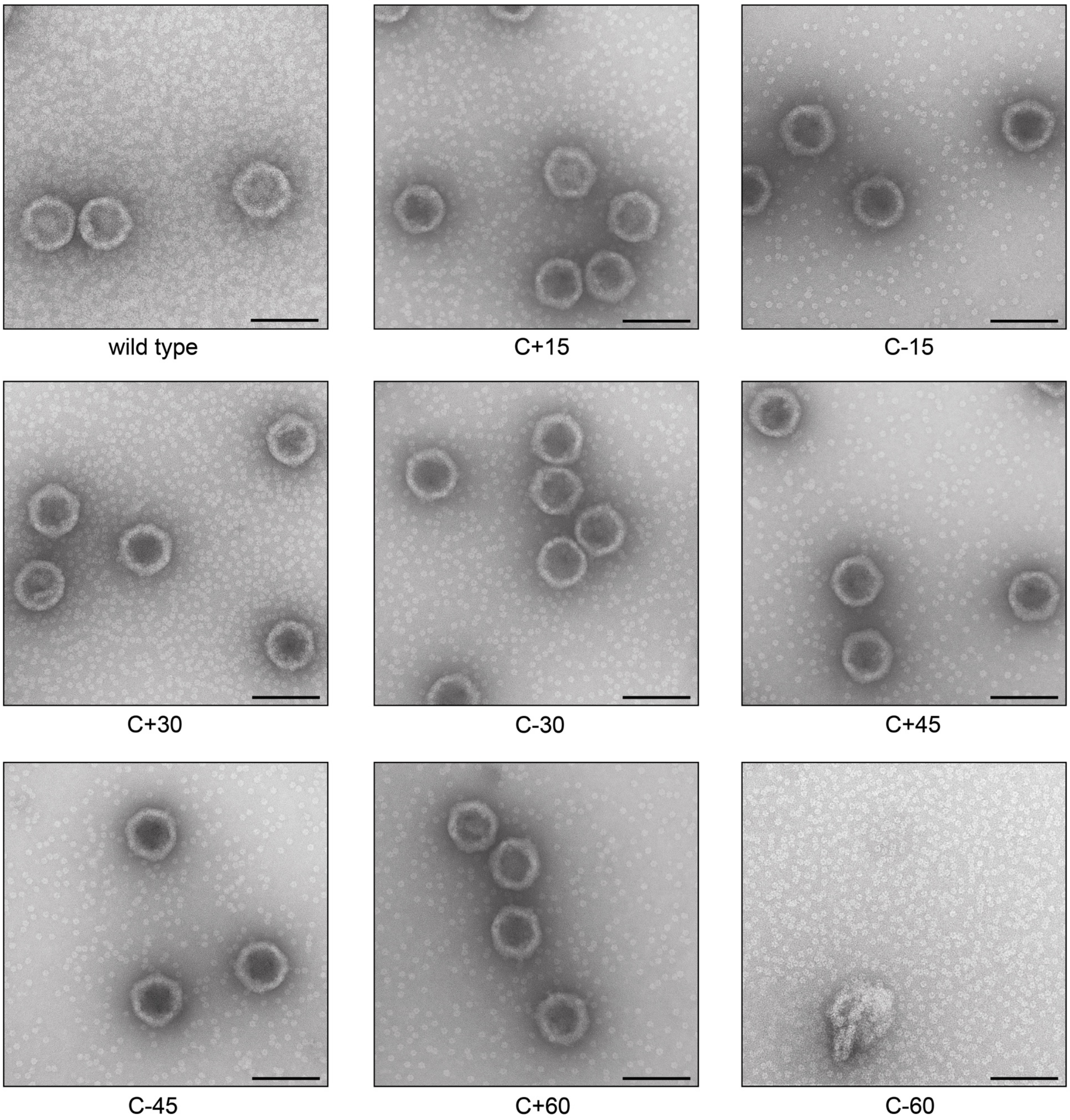


**Fig. S3. Dependence of capsid size on C-terminal length.** Negative-stain TEM of icosahedral-like whiffle balls assembled with 100 bp DNA in 60 mM UB-NaOH, 150 mM NaCl, pH 7.0. Assembly was carried out over 4 h with 1 mM $MCP_3$ and 30 ng/ml of the 100 pb dsDNA. The images represent typical Capsids of each sample with unassembled Capsomers in the background. No Capsids were assembled for the C-60 variant. Scale bars represent 100 nm.

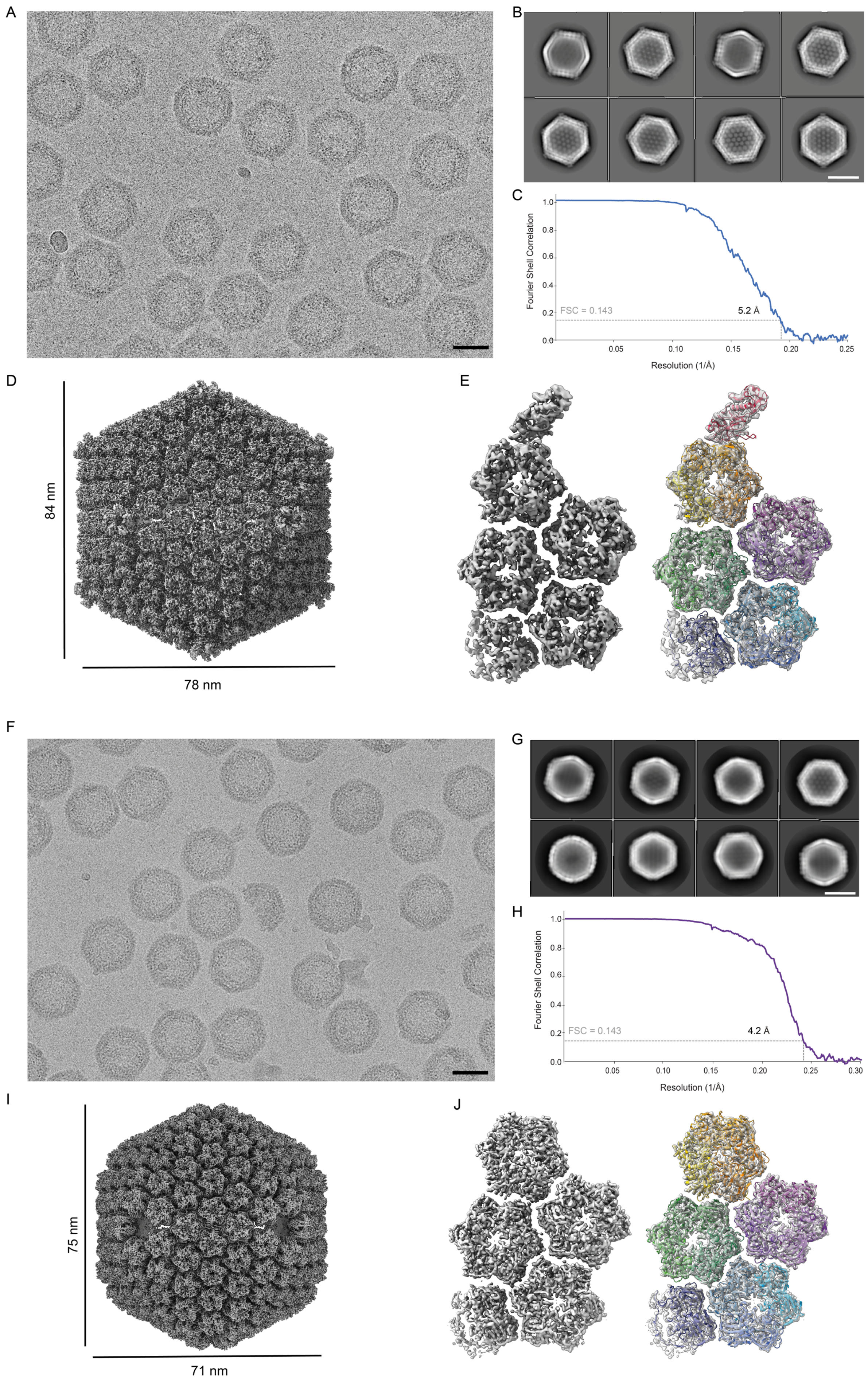
A
B
C
Fourier Shell Correlation
FSC = 0.143
5.2 Å
Resolution (1/Å)
D
84 nm
78 nm
E
F
G
H
Fourier Shell Correlation
FSC = 0.143
4.2 Å
Resolution (1/Å)
I
75 nm
71 nm
J

**Fig. S4. EM analysis of Mavirus VLP (A-E) and whiffle ball (F-J).** (**A**,**F**) Typical motion-corrected micrographs and (**B**,**G**) 2D class averages (scale bars, 50nm). (**C**,**H**) Fourier shell correlation (FSC) curves of the final, filtered density maps. The average resolution at 0.143 FSC criterion is indicated. (**D**,**I**) Complete cryo-EM density. (**E**,**J**) Cryo-EM density of the asymmetric unit (left) and with the fitted atomic model (right).

### Table 1. Cryo-EM data collection and model refinement statistics

| Data collection and processing | whiffle ball (MV18) | VLP (MV18+MV17) |
|---|---|---|
| Microscope | Titan Krios | |
| Voltage (kV) | 300 | 300 |
| Camera | K3 | Falcon 3 |
| Pixel size (Å) | 0.55 (super resolution mode) | 1.46 |
| Number of frames | 50 | 60 |
| Total electron dose ($e^-$/ Å $^2$) | 50 | 144 |
| Defocus range (µm) | 0.6 – 2.1 | 0.8 – 2.0 |
| Number of particles (initial / for final refinement) | 205098 / 47110 | 184255 / 70376 |
| Map resolution (Å) (FSC threshold) | 4.15 | 5.21 |
| Map sharpening B-factor (Å $^2$) | -210 | -271 |
| **R.m.s. deviations** | | |
| Bond length (Å) | 0.002 | 0.005 |
| Angles (°) | 0.485 | 0.626 |
| **Model validation** | | |
| Clashcore, all atoms | 5.25 | 10.09 |
| Poor rotamers (%) | 0.0 | 0.02 |
| Favored rotamers (%) | 99.7 | 99.56 |
| Ramachandran outliers (%) | 0.02 | 0.01 |
| Ramachandran favored (%) | 95.93 | 95.19 |
| Molprobity score | 1.65 | 1.86 |

### Nucleotide Sequences of final constructs of C-terminal variants

**MfeI_C1+15_HindIII**
CTGCGTAGTCCGCTGAAACTGACCGATGATAACAATAAACTGATGCGTGTGATGGGTGGTAGCTTT
ATGGGTGATGTTATGACCAACTTTAATCACATGGCAGCACATCCGGTTACCAAAACCGTGACCAAA
CTGCTGCGCAATGCAGGTCCTCTGAAAGACTATGCCGGTGATGGCACCATGATGGGTAATATTGCA
AGCGTTTATGGCTATGGCGGCAGTGGCGGCAGTGGCGGCAGTGGCGGCAGTGGCGGCAGTGGCAA
AAAAAAAACCACCACCCGGAAAAAAAAAAGGTGGTGAAATTGTTCTGCTGGGCAGCGGTAAAAAAG
GGGGTAAAAAACTGAGCGATAAACAGCTGCATGATCTGCGTAATCTGTAACTCGAGCGGCCGC

**MfeI_C1-15_HindIII**
CTGCGTAGTCCGCTGAAACTGACCGATGATAACAATAAACTGATGCGTGTGATGGGTGGTAGCTTT
ATGGGTGATGTTATGACCAACTTTAATCACATGGCAGCACATCCGGTTACCAAAACCGTGACCAAA
CTGCTGCGCAATGCAGGTCCTCTGAAAGACTATGCC↯GGCAAAAAAAAAACCACCACCCGGAAAA
AAAAAGGTGGTGAAATTGTTCTGCTGGGCAGCGGTAAAAAAGGGGGTAAAAAACTGAGCGATAAA
CAGCTGCATGATCTGCGTAATCTGTAACTCGAGCGGCCGC

**MfeI_C1+30_HindIII**
CTGCGTAGTCCGCTGAAACTGACCGATGATAACAATAAACTGATGCGTGTGATGGGTGGTAGCTTT
ATGGGTGATGTTATGACCAACTTTAATCACATGGCAGCACATCCGGTTACCAAAACCGTGACCAAA
CTGCTGCGCAATGCAGGTCCTCTGAAAGACTATGCCGGTGATGGCACCATGATGGGTAATATTGCA
AGCGTTTATGGCTATGGCGGCAGTGGCGGCAGTGGCGGCAGTGGCGGCAGTGGCGGCAGTGGCGG
CAGTGGCGGCAGTGGCGGCAGTGGCGGCAGTGGCGGCAGTGGCAAAAAAAAAACCACCACCCGG
AAAAAAAAAAGGTGGTGAAATTGTTCTGCTGGGCAGCGGTAAAAAAGGGGGTAAAAAACTGAGCG
ATAAACAGCTGCATGATCTGCGTAATCTGTAACTCGAGCGGCCGC

**Mfe_C1-30_HindIII**
CTGCGTAGTCCGCTGAAACTGACCGATGATAACAATAAACTGATGCGTGTGATGGGTGGTAGCTTT
ATGGGTGATGTTATGACCAACTTTAATCACATGGCAGCACATCCGGTTACCAAAACC↯GGCAAAA
AAAAAACCACCACCCGGAAAAAAAAAAGGTGGTGAAATTGTTCTGCTGGGCAGCGGTAAAAAAGGG
GGTAAAAAACTGAGCGATAAACAGCTGCATGATCTGCGTAATCTGTAACTCGAGCGGCCGC

**MfeI_C+45_HindIII**
CTGCGTAGTCCGCTGAAACTGACCGATGATAACAATAAACTGATGCGTGTGATGGGTGGTAGCTTT
ATGGGTGATGTTATGACCAACTTTAATCACATGGCAGCACATCCGGTTACCAAAACCGTGACCAAA
CTGCTGCGCAATGCAGGTCCTCTGAAAGACTATGCCGGTGATGGCACCATGATGGGTAATATTGCA
AGCGTTTATGGCTATGGCGGCAGTGGCGGCAGTGGCGGCAGTGGCGGCAGTGGCGGCAGTGGCGG
CAGTGGCGGCAGTGGCGGCAGTGGCGGCAGTGGCGGCAGTGGCGGCAGTGGCGGCAGTGGCGGCA
GTGGCGGCAGTGGCGGCAGTGGCAAAAAAAAAACCACCACCCGGAAAAAAAAAAGGTGGTGAAAT
TGTTCTGCTGGGCAGCGGTAAAAAAGGGGGTAAAAAACTGAGCGATAAACAGCTGCATGATCTGC
GTAATCTGTAACTCGAGCGGCCGC

**MfeI_C-45_HindIII**
CTGCGTAGTCCGCTGAAACTGACCGATGATAACAATAAACTGATGCGTGTGATGGGTGGTAGCTTT
ATGGGTGATGTT↯GGCAAAAAAAAAACCACCACCCGGAAAAAAAAAAGGTGGTGAAATTGTTCTGC
TGGGCAGCGGTAAAAAAGGGGGTAAAAAACTGAGCGATAAACAGCTGCATGATCTGCGTAATCTG
TAACTCGAGCGGCCGC

**MfeI_C+60_HindIII**
CTGCGTAGTCCGCTGAAACTGACCGATGATAACAATAAACTGATGCGTGTGATGGGTGGTAGCTTT
ATGGGTGATGTTATGACCAACTTTAATCACATGGCAGCACATCCGGTTACCAAAACCGTGACCAAA
CTGCTGCGCAATGCAGGTCCTCTGAAAGACTATGCCGGTGATGGCACCATGATGGGTAATATTGCA
AGCGTTTATGGCTATGGCGGCAGTGGCGGCAGTGGCGGCAGTGGCGGCAGTGGCGGCAGTGGCGG
CAGTGGCGGCAGTGGCGGCAGTGGCGGCAGTGGCGGCAGTGGCGGCAGTGGCGGCAGTGGCGGCA
GTGGCGGCAGTGGCGGCAGTGGCGGCAGTGGCGGCAGTGGCGGCAGTGGCGGCAGTGGCGGCAGT
GGCAAAAAAAAAACCACCACCCGGAAAAAAAAAAGGTGGTGAAATTGTTCTGCTGGGCAGCGGTAA
AAAAGGGGGTAAAAAACTGAGCGATAAACAGCTGCATGATCTGCGTAATCTGTAACTCGAGCGGC
CGC

**MfeI_C-60_HindIII**

CTGCGTAGTCCGCTGAAACTGACCGATGATAAC⚡GGCAAAAAAAAAAACCACCACCCGGAAAAAA
AAAGGTGGTGAAATTGTTCTGCTGGGCAGCGGTAAAAAAGGGGGTAAAAAACTGAGCGATAAACA
GCTGCATGATCTGCGTAATCTGTAACTCGAGCGGCCGC

The nucleotide sequences displayed here result in stepwise increases or decreases of 15 amino acid residues respectively where the increases are achieved through inserts of Gly-Gly-Ser repeats.
